# Mutations in *OsRZF1*, encoding a zinc-finger protein, causes reduced magnesium uptake in roots and translocation to shoots in rice

**DOI:** 10.1101/2022.06.02.494605

**Authors:** Natsuko I. Kobayashi, Hiroki Takagi, Xiaoyu Yang, Ayako Nishizawa-Yokoi, Tatsuaki Hoshina, Takayuki Oonishi, Hisashi Suzuki, Ren Iwata, Seiichi Toki, Tomoko M. Nakanishi, Keitaro Tanoi

**Affiliations:** Graduate School of Life Sciences, The University of Tokyo, 1-1-1 Yayoi, Bunkyo-ku, Tokyo 1138657, Japan; Faculty of Bioresources and Environmental Sciences, Ishikawa Prefectural University, 1-308 Suematsu, Nonoichi, Ishikawa, 921-8836, Japan; Institute of Agrobiological Sciences, National Agriculture and Food Research Organization (NARO), 3-1-3 Kannondai, Tsukuba, 305-8604, Japan; Center for Education and Research of Community Collaboration, Utsunomiya University, Utsunomiya 321-8505, Japan; National Institutes for Quantum and Radiological Science and Technology, 4-9-1 Anagawa, Inage-ku, Chiba, 263-8555, Japan; Cyclotron and Radioisotope Center (CYRIC), Tohoku University, 6-3 Aramaki Aza-Aoba, Aoba-ku, Sendai, 980-8572, Japan; Faculty of Agriculture, Ryukoku University, 1-5 Yokotani, Seta Oe-cho, Otsu, Shiga 520-2194, Japan; Graduate School of Nanobioscience, Yokohama City University, 22-2 Seto, Yokohama, Kanagawa, 236-0027, Japan; Hoshi University, 2-4-41 Ebara, Shinagawa-Ku, Tokyo, 142-8501, Japan

## Abstract

Magnesium (Mg) homeostasis is critical for maintaining many biological processes, but little information is available to comprehend the molecular mechanisms regulating Mg concentration in rice *(Oryza sativa).* To make up for the lack of information, we aimed to identify mutants defective in Mg homeostasis through a forward genetic approach. As a result of the screening of about 3,000 M2 seedlings mutated by ion-beam irradiation, we found a rice mutant that showed reduced Mg content in leaves and slightly increased Mg content in roots. Radiotracer ^28^Mg experiments showed that this mutant, named low magnesium content 1 (LMGC1), has decreased Mg^2+^ influx in the root and Mg^2+^ translocation from root to shoot. The MutMap method identified 7.4 kbp deletion in the LMGC1 genome leading to a loss of two genes. Genome editing using CRISPR-Cas9 further revealed that one of the two lost genes, a gene belonging to RanBP2-type zinc finger family, was the causal gene of the low-Mg phenotype. Considering this gene, named *OsRZF1,* has never been reported to be involved in ion transport, the phenotype of LMGC1 would be associated with a novel mechanism of Mg homeostasis in plants.

## Introduction

Magnesium (Mg) is the second most abundant intracellular cation in plant cells. It exists predominantly as a complex form with cellular components with a unique binding-affinity for each (Cowan, 2002), and the concentration of free Mg^2+^ in the cytosol is about 0.3 mM (Maguire and Cowan, 2002; Gout et al., 2014). In the cellular components, Mg is efficiently incorporated with various metabolic cycles by serving a catalytic role and allosteric effects (Pierce and Reddy, 1986).

In response to the limited Mg availability in the environment, the decrease in the phloem loading of sucrose is detected as one of the signs because H^+^-ATPases with consuming ATP energy is sensitive for Mg deficiency, and the proton-motive force created by H^+^-ATPases drives sucrose transporters that mediate phloem sucrose loading (Geiger, 2011). After that, the overaccumulation of photoassimilates, and finally onset of leaf interveinal chlorosis, have been observed in this order in bean (Cakmak et al., 1994), sugar beet (Hermans et al., 2004), and *Arabidopsis* (Hermans and Verbruggen 2005; Ogura et al., 2020). Characteristics of Mg deficiency compared to other mineral deficiencies can be the high susceptibility of young mature leaves. Whole plant iodine staining revealed that starch accumulates preferentially in the most recently expanded young leaves (Hermans et al., 2004; Hermans and Verbruggen 2005; Kobayashi et al., 2013; Ogura et al., 2020). These leaves can already function as the source with increasing photosynthetic activity while the Mg concentration drastically decreases during leaf expansion. The Mg concentration below 1-2 mg g^-1^ can be the benchmark for the onset of chlorosis in any plant species (Hermans et al., 2013). At about the same time or later than the chlorosis appears, leaf growth rate started to be reduced (Hermans and Verbruggen 2005) most probably due to the shrinking carbon partitioning and photosynthesis. The degree of the growth retardation and the time of its appearance will vary depending on the age of the plant materials used in the experiment (Hermans et al., 2013), as they can relate to Mg storage amounts and Mg retranslocation capability in plant tissues.

Magnesium is known to be mobile in many plant species (Hocking, 1994; Maillard et al., 2015). In rice plants, the concentration of Mg^2+^ in phloem sap was reported to be 12 mM (Fukumorita et al., 1983), which is apparently higher than the Mg^2+^ concentration in the cytosol. For the active Mg retranslocation, removal of Mg from chlorophyll structure by Mg dechelatase SGR is involved (Peng et al., 2019), but other details of the retranslocation process including phloem Mg^2+^ loading are not yet known. Magnesium uptake and xylem transport is also properly controlled. The concentration of Mg^2+^ in rice xylem sap is concentrated compared to the external solution when the Mg^2+^ concentration of the external solution is less than 3 mM (Tanoi et al., 2011). In Arabidopsis, MGR4 and MGR6 are responsible in this mechanism as xylem loaders in the root stele (Meng et al., 2022) and MTP10 acts as a xylem unloader in the shoot parenchyma cells in vascular bundles (Ge et al., 2022). Mutations in these transporters cause modification in Mg content in the whole plant. For the Mg^2+^ uptake system, AtMRS2-4/MGT6 (Mao et al., 2014), OsMGT1/OsMRS2-2 (Chen et al., 2017), and OsCZMT1 (Lim et al., 2022) are suggested to be involved, although the mechanistic basis for their function has not fully been elucidated. Keeping magnesium stored in the vacuole is another important strategy for surviving environmental fluctuations. AtMRS2-1/MGT2 (Conn et al., 2011), AtMRS2-5/MGT3 (Conn et al., 2011), and MGR1 (Tang et al., 2022) are the tonoplast localized transporters participating in this process to confer Mg excess stress. Loss of these molecules results in decreased Mg content in leaves. Apart from the ion transporters, the root endodermal barrier contributes Mg homeostasis in leaves, given that the mutation in SHENGEN3 leads to Mg overaccumulation in *Arabidopsis* (Pfister et al., 2014) and *Brassica rapa* (Alcock et al., 2021).

These previous works are gradually revealing the molecules that act as valves on Mg ion distribution circuit. Now, it is necessary to identify more components involved in the Mg homeostasis to understand the whole picture. In this study, we physiologically and genetically characterized a rice mutant with low Mg content in leaves, which we have isolated from a screening of ion-beam mutagenized M2 population. The mutant, named low magnesium content 1 (LMGC1), showed reduced Mg^2+^ influx and impaired root-to-shoot Mg transport from the early stages of growth. The causative mutation of the low Mg phenotype of LMGC1 was found by MutMap analysis, leading to the identification of the causal gene whose involvement in Mg homeostasis had never been considered before.

## Results

### Disturbance in Mg distribution in LMGC1

LMGC1 is the mutant originally selected by the decreased Mg content in leaves under low Mg condition. Consistent with the original observation, Mg content of the shoot in LMGC1 grown with the nutrient solution for one week was decreased compared to the wild-type regardless of the Mg^2+^ concentration in the nutrient solution (Fig. 1A). The largest decrease was 35%, which was observed under the control condition (Fig. 1A). In the root, however, LMGC1 was found to accumulate more Mg than the wild-type under control condition and low Mg condition (Fig. 1B). As a result, the distribution of Mg to the shoot and the root was greatly altered in LMGC1.

**Figure 1.**
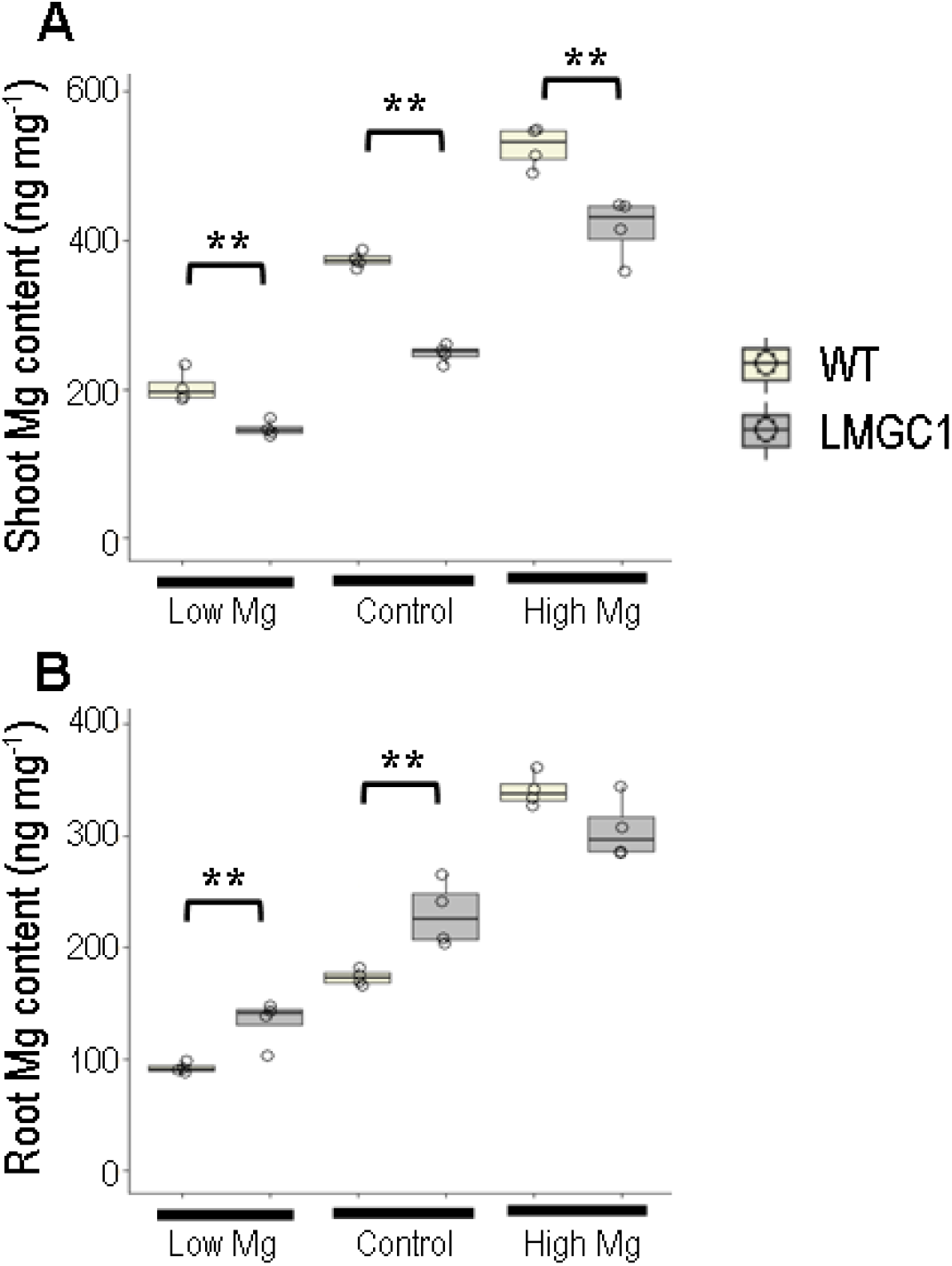
Magnesium content in shoots (A) and roots (B) of wild-type and LMGC1 mutant (n = 4). Following the cultivation in 0.5 mM CaCl_2_ solution, rice plants were grown for 1 week in the nutrient solution containing either 27 μM (low Mg), 0.27 mM (control), or 2.27 mM (high Mg) of Mg^2+^. Asterisks indicate significant differences between the wild-type (WT) and LMGC1 mutant (\*\**p* < 0.01, Student’s *t*-test).

### Growth retardation observed under low Mg condition

The effect of the low-Mg content in shoot on the plant growth was examined. After 22 days, LMGC1 grew similarly as the wild-type unless Mg was supplied enough (Fig. 2A). Under low Mg condition, the color was getting lighter from the tips of the young mature leaves, and the leaves of LMGC1 were further beginning to wither (Fig. 2A). Accordingly, SPAD value in the 5th leaf, which was one of the mature leaves, was lowered (Fig. 2B), and the shoot biomass was decreased (Fig. 2C) in LMGC1 compared to the wild-type under low Mg condition. In contrast, root growth was not reduced significantly in LMGC1 (Fig. 2D).

**Figure 2.**
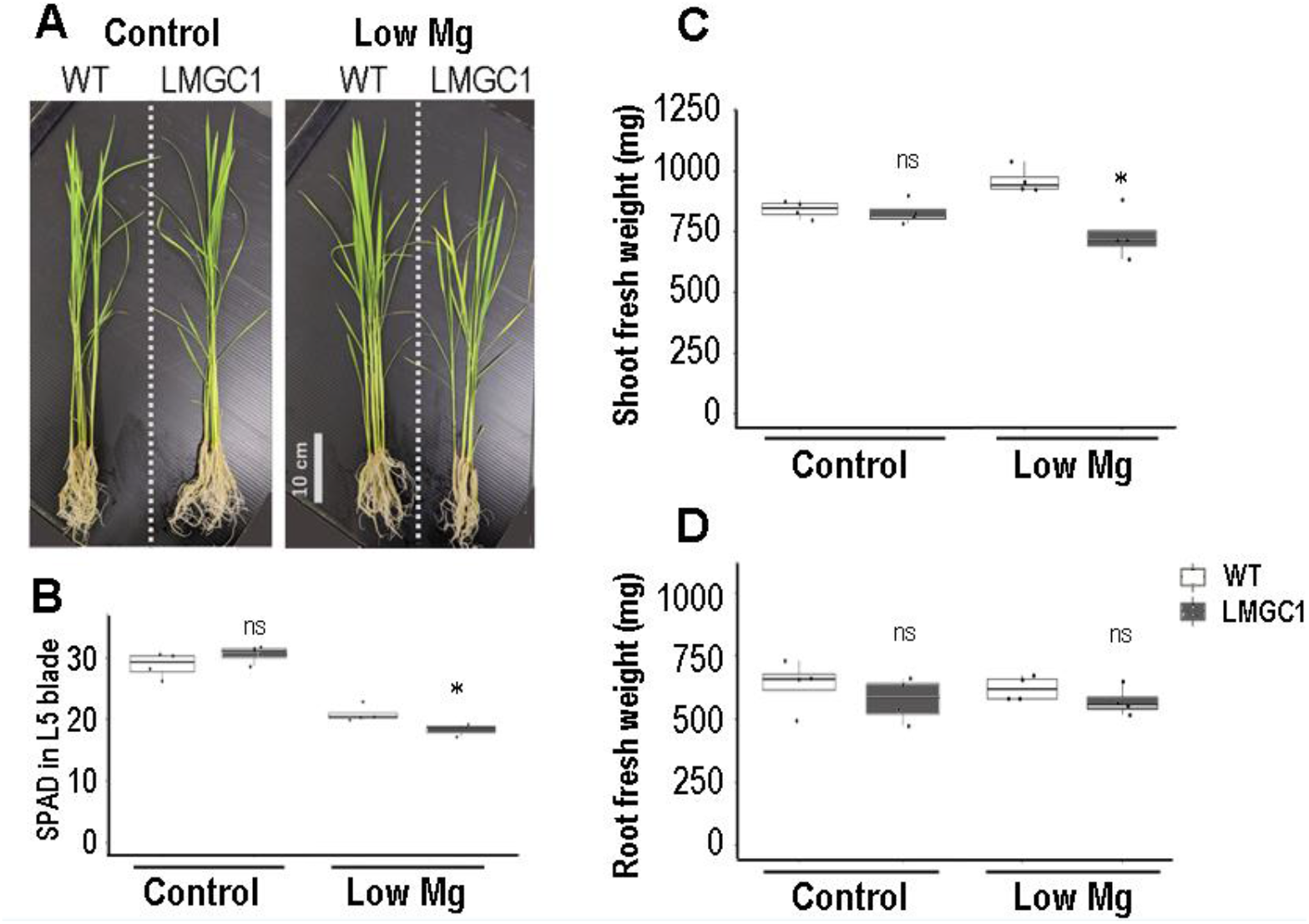
Phenotypes of LMGC1 after cultivation in the nutrient solution containing either 14 μM (low Mg) or 0.27 mM (control) of Mg^2+^ for 22 days. (A) Changes in appearance (n = 4). (B) SPAD value at the leaf blade of the 5th leaf (n = 3-4). (C) The fresh weight of the shoot (n = 4). (D) the fresh weight of the root (n = 4). “ns” indicates not significant *(p* > 0.05), whereas the asterisk indicates significant difference *(p* < 0.05) between the wild-type and LMGC1 mutant (Student’s *t*-test).

### Magnesium uptake kinetics and translocation from roots to shoots

To understand how Mg distribution in LMGC1 altered, the Mg^2+^ dynamics in the seedlings grown under control condition for 1 week was analyzed using the radiotracer ^28^Mg. The Mg^2+^ uptake rate in LMGC1 was found to be reduced by half in any Mg^2+^ concentration in the uptake solution (Fig. 3A). Focusing on the experiment with 0.25 mM Mg^2+^ concentration in the uptake solution, the shoot-to-root ratio of the ^28^Mg was less than one-third for LMGC1 compared to the wild-type (Fig. 3B).

**Figure 3.**
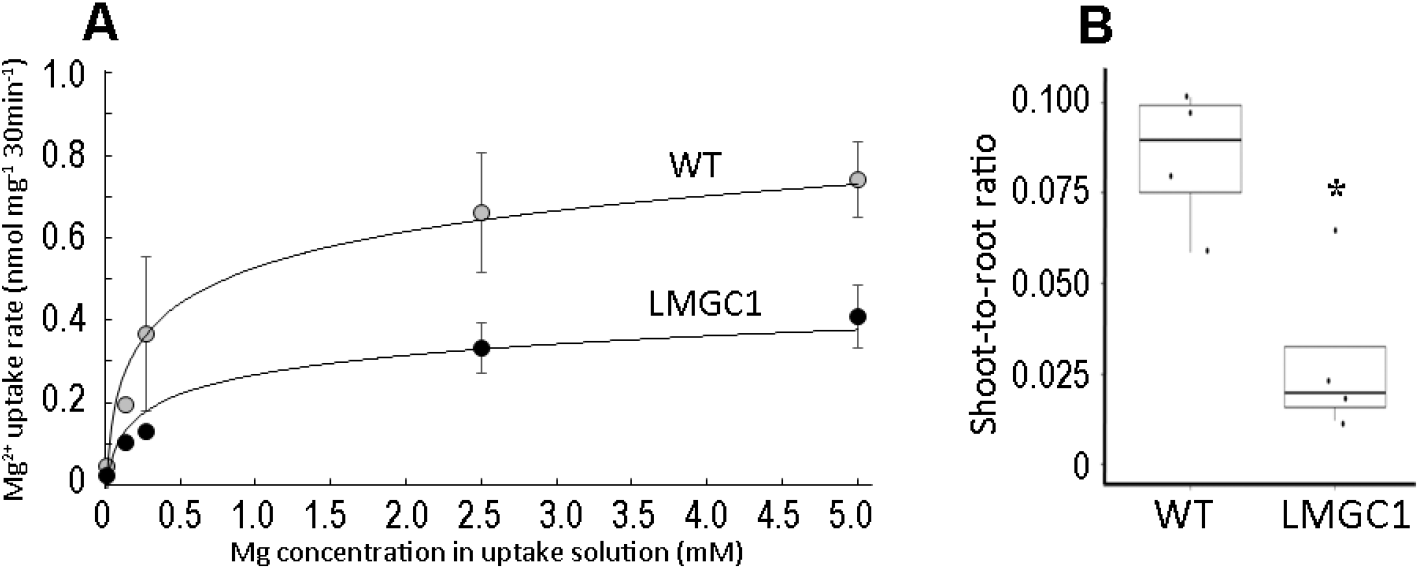
The kinetic analysis of Mg^2+^ uptake and transport in the wild-type (WT) and LMGC1 plants grown in the nutrient solution (0.27 mM Mg^2+^) for 1 week following the cultivation in CaCl_2_ solution. (A) The Mg^2+^ uptake rate determined in the nutrient solution with various Mg^2+^ concentration (15-5000 μM) (n = 4). (B) The shoot-to-root ratio in the ^28^Mg distribution in the uptake experiment using 0.25 mM Mg^2+^ solution (n = 4) (\**p* < 0.05, Student’s *t*-test).

### Modified Mg^2+^ dynamics in LMGC1 found the day after transplanting to the nutrient solution

With the result that the amount of Mg^2+^ uptake in 30 minutes was reduced in LMGC1, we tried to distinguish whether the influx had decreased, or the efflux had increased. Based on the ^28^Mg signal intensity after 20 seconds’ absorption, the Mg^2+^ influx rate (nmol mg^-1^ min^-1^) was calculated to be 0.126 for wild-type and 0.088 for LMGC1 (Fig. 4A). The distribution image of ^28^Mg in the seedlings showed that the signal intensity in LMGC1 was reduced throughout the root, but the signal distribution was not different from the wild-type (Fig. 4B). The efflux rate (nmol mg^-1^ min^-1^) was calculated to be 0.052 for wild-type, and 0.050 for LMGC1 based on the y-intercept of the linear regression (Fig. 4C, 4D). The experiment was repeated twice, and the results of the second experiment were 0.051 and 0.060 for wild-type and LMGC1, respectively. The difference in efflux rate between the wild-type and LMGC1, if any, was found to be obviously smaller than the difference in the influx rate.

**Figure 4.**
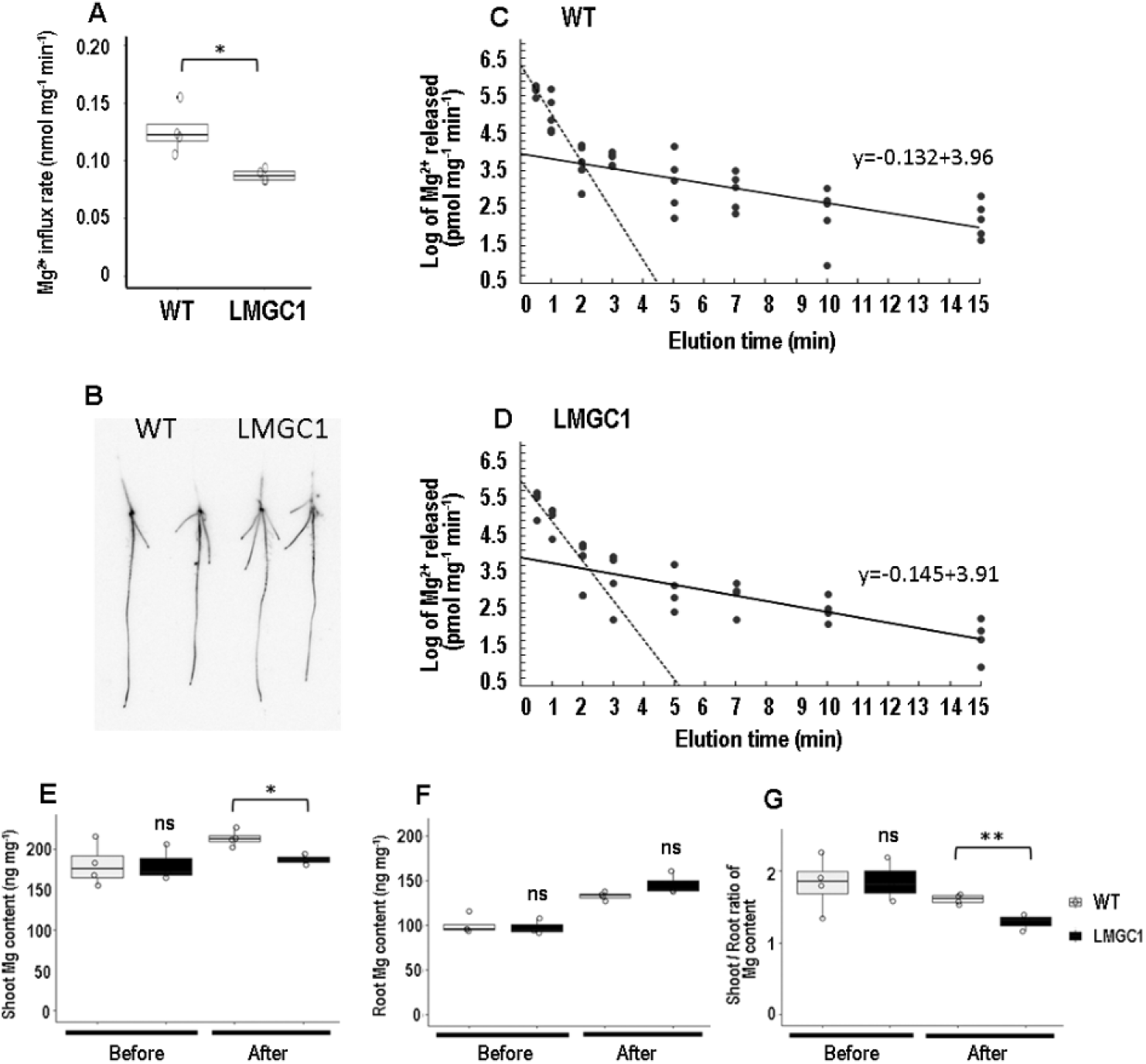
Magnesium ion flux and distribution in the wild-type (WT) and LMGC1 mutants cultivated in the nutrient solution (0.27 mM Mg^2+^) for 1 day following the cultivation in CaCl_2_ solution. (A) Magnesium influx determined based on the ^28^Mg radioactivity after 20 seconds’ absorption (n = 4). (B) The ^28^Mg radioactivity image in the seedlings after 20 seconds’ absorption (n = 2). (C-D) Time-course of the Mg^2+^ elution in the wild-type (C) and LMCG1 mutant (D). Semi-logarithmic plots were made using 4-5 seedlings to construct the efflux curves. The efflux rate (nmol mg^-1^ min^-1^) was determined based on the y-intercept of the linear regression. The experiments were conducted twice, and the first experimental results were presented. (E-G) Magnesium content in the seedlings before transplantation to the nutrient solution (0.27 mM Mg^2+^) and 24 h after transplantation (n = 4). (G) The shoot-to-root ratio of Mg content. Asterisks indicate significant differences between the wild-type and LMGC1 mutant (\*\**p* < 0.01, \**p* < 0.05), whereas “ns” indicates there is no significant difference (*p* > 0.05) (Student’s *t*-test).

Given that the decrease in the influx rate in LMGC1 was detected as early as the day after transplanting into the nutrient solution, we proceeded to test the possibility that the decrease in influx rate preceded the disturbance in Mg distribution. The Mg content in the shoot of LMGC1 was comparable to that of the wild-type before transplanting to the nutrient solution, but it was lower than the wild-type the day after transplanting into the nutrient solution (Fig. 4E). In the root, although there was no statistically significant difference, Mg content of LMGC1 tended to increase compared to the wild-type after transplanting to the nutrient solution (Fig. 4F). Consequently, the shoot/root ratio of Mg content was significantly lower in LMGC1 than the wild-type already the day after transplanting to the nutrient solution (Fig. 4G).

### The large deletion in the chromosome 1

First, F2 were developed by a cross LMGC1 with wild type Koshihikari. Among 90 individuals of the F2, 10 individuals containing highest and low concentration of Mg were applied to MutMap+ approach as high and low bulk samples, respectively. A genomic region having contrast SNP-index value (Abe et al. 2012) between high and low bulk samples was identified on at 20 Mb on chromosome 1 (Fig. 5A). However, no candidate SNP causing amino acid change were detected. Next, therefore, we compared the sequence depth between bulk samples around 20 Mb on chromosome 1 for identifying the candidate mutation caused by changing genomic structure. As a result, 7.4 kb at 20,932,901 – 20,940,302 on chromosome 1 had few short reads in specifically low Mg bulk samples (Fig. 5B), implying that this 7.4 kb should be candidate deletion for low Mg concentration in LMGC1. By this 7.4 kb deletion, Os01g0555100 and Os01g0555200 are completely and partially excluded from the genome (Fig. 5C). To verify that the 7.4 kb deletion is associated with low Mg phenotype of LMGC1, we confirmed the association between leaf Mg content and genotype in 80 individuals of F2 progeny derived from a different F1 individual from the one used for MutMap+ analysis. All 17 individuals with homozygous 7.4 kb deletion confirmed by PCR had low Mg content compared with the individuals having WT genotype in homozygous and heterozygous, indicating that this deletion should be highly candidate for causal mutation for low Mg phenotype of LMGC1 (Fig. 5D).

**Figure 5.**
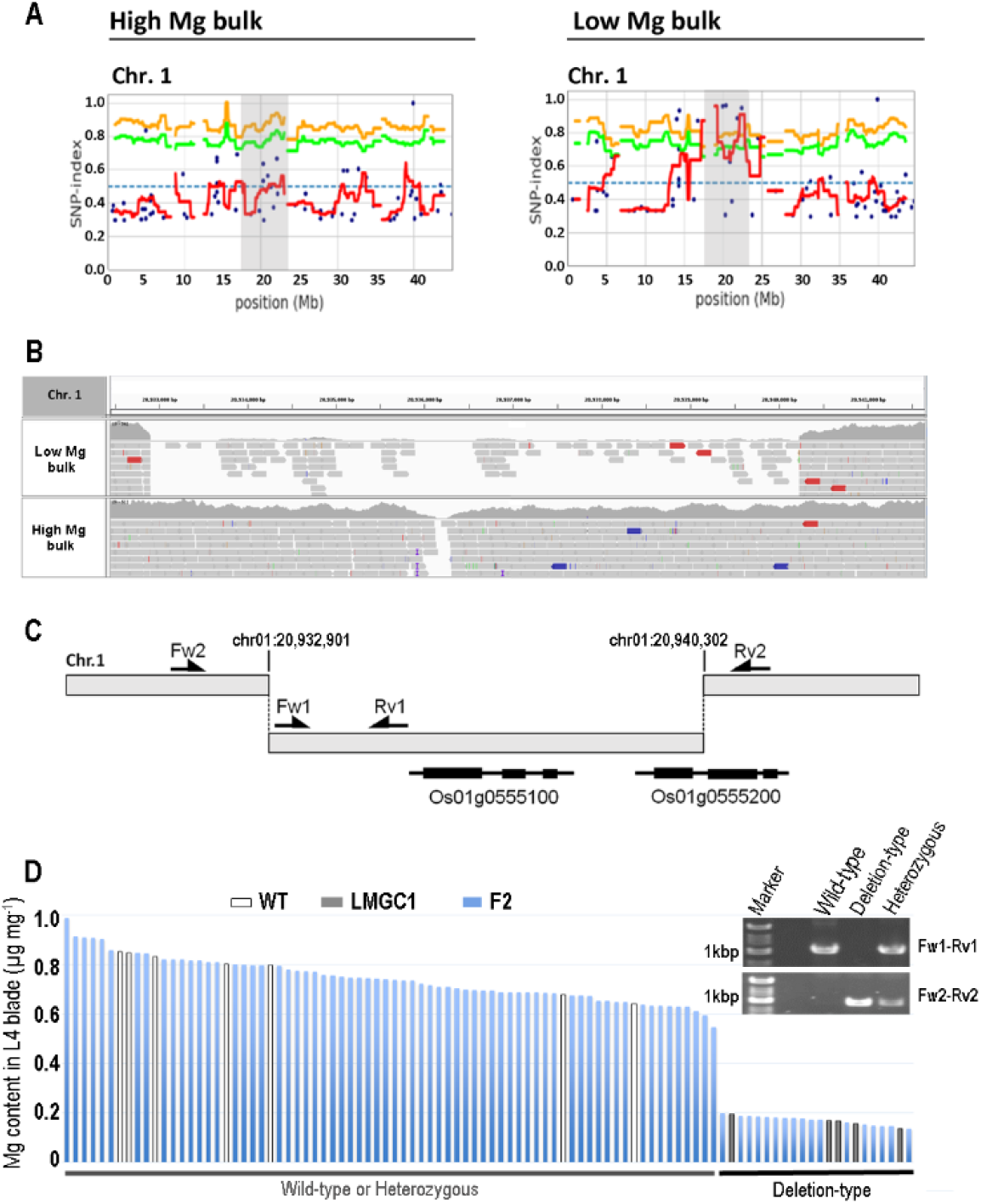
Identifying the causal mutation for low Mg content of LMGC1. (A) The identification of a candidate genomic region (gray) on chromosome 1 based on MutMap+ analysis. The blue dots represent a SNP-index value at a given SNP position. Red line represents the sliding window average SNP-index values of 2 Mb interval with 50 kb increment. The green and yellow lines are the 95% and 99% confidence interval of SNP-index value under the null hypothesis of a randomly bulked DNA sample, respectively. (B) The visualization of the aligned sequence reads around 20Mb on chromosome 1 with Integrative Genomics Viewer. The region covered by pink shadow showed the region aligning few sequence reads in only low bulk sample. (C) The schematic illustration of the physical relationship between the large deletion and the localizing tow genes, *Os01g0555100* and *Os01g0555200.* The primer used for deciding genotype in the large deletion were illustrated in allows (Fw1-Rv1 and Fw2-Rv2). (D) The relationship between the leaf Mg content and genotype in the F2 progenies. The primer pairs, Fw1-Rv1 and Fw2-Rv2, amplifies about 1 kb PCR product in wild type allele and 0.9 kb mutation allele from large deletion, respectively. The individuals showing PCR products from both primer pair indicates heterozygous genotype in this locus.

### Identification of the causal gene for LMGC1

The first candidate is Os01g0555100, previously identified as one of the stress repressive zinc finger proteins (Huang et al., 2008) belonging to the RanBP2-type zinc finger protein family (RZF). For the second candidate of Os01g0555200 gene, predicted amino acids sequences indicate a Asp/Glu racemase family protein (Rac). To validate the causal gene for LMGC1, we generated mutants using CRISPR/Cas9-targeted mutagenesis in the Nipponbare rice cultivar, and investigated the Mg content in the leaf. The mutants RZF #6-3 and RZF #8-3 had deletions of 54 bp and 38 bp in Os01g0555100, respectively (Fig. 6A). Both deletion mutations resulted in the loss of one of the three zinc finger domains, and the 38 bp deletion further caused the frameshift. Rac #23-1 was a mutant with 126 bp deletion, and Rac #23-2 had a 3 bp plus 13 bp deletions (Fig. 6B). As a result of the 126 bp deletion, 42 residues out of a total of 329 residues in Rac protein were lost. The mutation identified in Rac #23-2 leads to a significant alteration in the amino acids sequence including a loss of the asparagine and threonine residues functioning as a binding site for the substrate (Ahn et al., 2015). Measurement of Mg content in the leaf of these mutant lines revealed that mutations in RZF gene, but not Rac gene, can reproduce the low Mg phenotype of LMGC1 (Fig. 6C). We named this gene *OsRZF1* and performed expression analysis to explore its potential role. The gene was found to accumulate in both leaves and roots, with more in the leaves (Fig. 6D). Transient expression assay using tobacco leaf epidermis showed EGFP fluorescence not localized to a specific subcellular compartment (Fig. 6E), indicating that OsRZF1 is a cytosolic protein.

**Figure 6.**
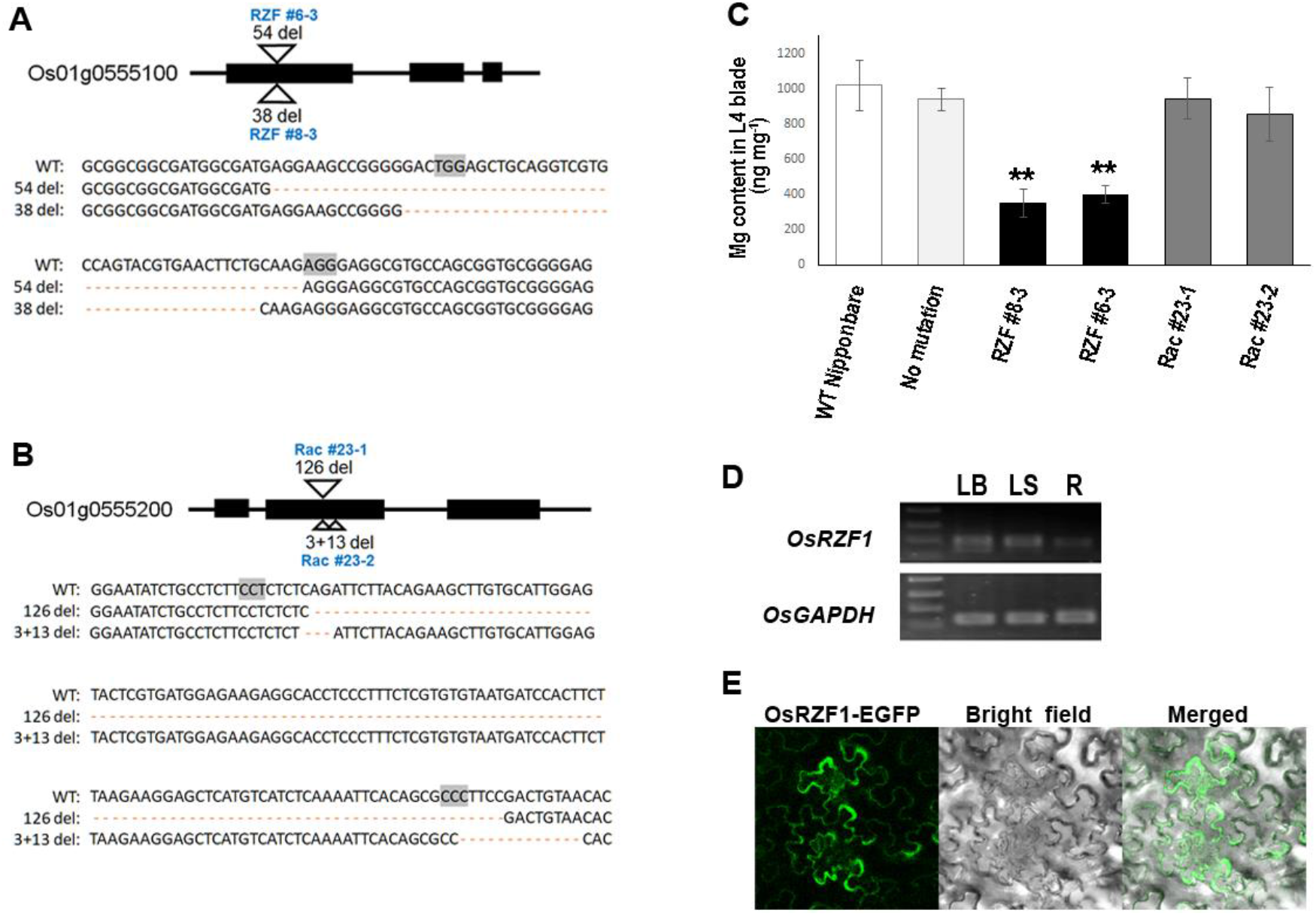
Identification of the *OsRZF1* through the analysis of CRISPR/Cas9-mediated mutant lines, and determination of its expression and localization. (A) RZF #6-3 and RZF #8-3 have the deletion mutations in the first exon of Os01g0555100 gene. A part of the coding sequence (13 – 111 bp in the WT) is presented. (B) Rac #23-1 and Rac #23-2 have the deletion mutations in the second exon of Os01g0555200 gene. A part of the coding sequence (277 – 437 bp in the WT) is presented. The PAM sequences used for the genome editing were colored in gray. (C) Magnesium content in the blade of the 4th leaf of the mutant lines. Wild-type Nipponbare plant and the agrobacterium-infected plant which gained antibiotic resistance but failed to cause any mutation in the targeted gene (no mutation) were used as the reference (n = 4-7). The mutants with asterisks are significantly different from the wild-type Nipponbare (\*\**p* < 0.01, Student’s *t*-test). (D) Expression of *OsRZF1* gene analyzed by RT-PCR. *OsGAPDH* was used as a reference. LB; leaf blade, LS; leaf sheath, R; root. (E) Transient expression assay of OsRZF1-EGFP fusion protein in *Nicotiana benthamiana* leaf epidermal cells.

## Discussion

Using ^28^Mg, a very rare material, it was found that the reduced influx (Fig. 4A) and unchanged efflux (Fig. 4C, 4D) achieved the lowered Mg uptake in LMGC1 (Fig. 3A). The process of this experiment also yielded the quantitative relationship between the influx and efflux to uptake of Mg^2+^ in the rice root. The calculated efflux rate in the wild-type was 0.052 nmol mg^-1^ min^-1^ (Fig. 4C), and the difference between this and the influx rate, 0.126 nmol mg^-1^ min^-1^, corresponded to the uptake rate, namely, 0.074 nmol mg^-1^ min^-1^. It is suggested that Mg homeostasis in rice roots is maintained by a futile cycling system in which nearly half of the Mg^2+^ entering the root cells is discharged and only the other half is acquired. This idea is comparable with the very close values of Mg^2+^ efflux and influx in excised onion roots (Macklon and Sim, 1976).

In ion uptake and translocation, the main molecules may be distinct, or there may be some molecules involved in both. In the former case, the ion uptake from the rhizosphere into the root outer cells can be mediated by the influx transporters such as potassium transporter HAK5 (Rubio et al., 2000; Qi et al., 2008) and manganese transporter OsNramp5 (Sasaki et al., 2012), whereas xylem loading is implemented by the efflux transporters functioning in ion exclusion from cells in the neighborhood of xylem such as potassium SKOR channel (Gaymard et al., 1998), H^+^/K^+^ antiporter NRT1;5 (Li et al., 2017), and manganese transporter OsMTP9 (Ueno et al., 2015). The lack of any one of these transporters significantly reduces the above-ground content of the ions that the transporter was carrying (Gaymard et al., 1998; Nieves-Cordones et al., 2010; Sasaki et al., 2012; Ueno et al., 2015). In case of Mg^2+^, the fact that xylem Mg^2+^ loading is mediated by the efflux-type transporters in *Arabidopsis* (Meng et al., 2022) suggests the systems for uptake and translocation are distinct. We thought that if it was the Mg^2+^ influx function that was directly impaired in LMGC1, and the reduction in Mg^2+^ transport to the shoot was a secondary effect, then the reduction in influx would have occurred prior to the reduction in Mg^2+^ translocation. However, in the present experiment, both phenotypes were observed just the day after transplanting into the nutrient hydroponic solution (Fig. 4A, 4G). Therefore, it is impossible to determine at present whether one of the two phenotypes; Mg^2+^ influx and translocation, is the primary cause or whether both occur independently downstream of OsRZF1.

Investigations on CRISPR/Cas9-mediated mutants demonstrated that the Mg homeostasis in rice requires *OsRZF1* gene, expressed in the whole plant. The protein encoded by *OsRZF1* will contain three RanBP2-type zinc finger domains. The consensus sequence of this domain is W-X-C-X_2-4_-C-X_3_-N-X_6_-C-X_2_-C (C4-type), which recognizes ssRNA (Loughlin et al., 2009; Nguyen et al., 2011). In plants RZF phylogenetic tree, *OsRZF1* is in the same clade as rice *stress repressive zinc finger 1 (SRZ1)* (Huang et al., 2008) and *Arabidopsis stress associated RNA-binding protein 1 (SRP1)* (Xu et al., 2017). Both *SRZ1* and *SRP1* expressions are responsive to ABA, salt, and cold treatments (Huang et al., 2008; Xu et al., 2017). In the ABA signaling process, SRP1 acts as a positive regulator having the ability to directly bind to 3’UTR mRNA of *ABI2* to reduce its stability (Xu et al., 2017). Hence, some role in regulating gene turnover and /or gene stability has been predicted also for other proteins belonging to this clade (Gipson et al., 2020). Nevertheless, in consideration that SRP1 localizes in the nucleus while OsRZF1 is potentially the cytosolic protein, their mode of function might be different. In studies of RZF proteins, the connection to elemental transport has never been investigated, and the results of this study are the first to demonstrate a fundamental role for RZF in mineral homeostasis.

## Materials and Methods

### Plant growth conditions

Rice plants were germinated in pure water and grown further in 0.5 mM CaCl_2_ solution (Ca solution) for 2-3 days, then transferred to the half strength Kimura B nutrient solution (Kobayashi et al., 2018). Magnesium concentration in the nutrient solution was modified by adding MgSO_4_ or substituting MgSO_4_ for Na_2_SO_4_ according to the experimental subjects. The nutrient solutions were refreshed every 2-3 days. Plant incubator was set to 28 °C with 10:14-h light/dark photoperiod.

### Elemental analysis

Rice *(Oryza sativa.* L. Koshihikari) wild-type and LMGC1 mutant plants grown for 1 week in the nutrient solution containing either 27 μM (low Mg), 0.27 mM (control), or 2.27 mM (high Mg) of Mg^2+^ were harvested. Rice seedlings were also sampled just before the transplanting into the nutrient solution and one day after transplanting. Nipponbare cultivar wild-type and genome-edited mutant lines generated by the method shown in the later section were also used for elemental analysis. They were grown in the control nutrient solution for 2 weeks and the blade of the 4th leaf was sampled. Shoots, roots, or leaf blades were separately weighed and digested completely in 69% nitric acids at 90 °C by a closed vessel digestion procedure. Sample solutions were analyzed with Inductively Coupled Plasma-Mass Spectrometry (NexION, PerkinElmer) after being diluted appropriately with ultrapure water. The quantification of the method was verified using the certificated reference material (NCS DC73349, China National Analysis Center).

### Growth assay

Wild-type and LMGC1 mutant plants were grown under either 14 μM Mg (low Mg) or 0.27 mM Mg (control) conditions for 22 days until the 7th leaf was matured. Then, the SPAD value at the leaf blade of the 5th leaf was determined using a chlorophyll meter (SPAD-502 plus, Konica Minolta), and the fresh weight of shoots and roots was measured.

### Magnesium uptake, translocation, and flux analysis using radiotracer

Carrier-free ^28^Mg with a half-life of 21 h was produced by the ^27^Al(α,3p)^28^Mg reaction in the cyclotron and chemically purified (Tanoi et al., 2013). Wild-type and LMGC1 mutant plants were grown in the nutrient solution containing Mg^2+^ of 0.27 mM for 1 week. For the measurement of Mg uptake rate, the roots of the plants were soaked for 30 min in the uptake solution, that was the half strength Kimura B nutrient solution with any Mg concentrations (15-5000 μM) labeled with ^28^Mg. After absorption, the roots were rinsed with ice-cold nutrient solution for 10 min. Then, the remaining solution on the surface was wiped off with a paper towel, and the samples were separated into the shoot and the root, weighed, and subjected to the gamma-ray counting (Kobayashi et al., 2018). The amount of Mg accumulated in the shoots and the roots during 30 min was added up and divided by the root weight to obtain Mg uptake rate (nmol mg^-1^ 30 min^-1^). Additionally, the translocation of Mg^2+^ from root to shoot was calculated based on the ^28^Mg concentration ratio of the shoots and the roots.

For the Mg^2+^ flux analysis, Mg^2+^ influx and efflux in the root were independently measured. The roots of the plants grown with the nutrient solution with 0.27 mM Mg for 24 h were soaked in the uptake solution containing 0.27 mM Mg labeled with ^28^Mg (4.8 kBq/ml) for 20 seconds, and then rinsed for 30 seconds in ice-cold nutrient solution. Weighing and the radioactivity measurement were performed as described above. Based on the Mg amount absorbed in 20 seconds, the influx rate was determined to be the value converted to the amount absorbed per minute per unit weight of the root (nmol mg^-1^ min^-1^). In addition, the distribution of ^28^Mg in the seedlings after being rinsed was visualized using the imaging plate and image reader (Kobayashi et al., 2013).

To analyze the efflux rate, the seedlings grown in the CaCl_2_ solution were transferred to the same uptake solution that was used in the influx analysis, and left for labelling for 24 h to ensure that all Mg^2+^ taken into the roots from the nutrient solution was uniformly labeled with ^28^Mg. After labelling, the roots of the plants were quickly dipped in the non-labeled nutrient solution to remove the surface-remained ^28^Mg, and then soaked in the series of non-labeled nutrient solution for elution of ^28^Mg for 0.5, 0.5, 1, 1, 2, 2, 3, and 5 min. The total duration of elution was 15 min. The amount of Mg^2+^ released from the root per minute per unit weight of the root (pmol mg^-1^ min^-1^) was determined based on the radioactivity in the elution solution and the assumption that the specific radioactivity of Mg^2+^ released from the roots was equal to the uptake solution (Pitman, 1971). The basic theory to determine the efflux rate was that of compartmental analysis, established by the 1990s (Lee and Clarkson 1986; Kronzucker et al., 1995). The amount of Mg^2+^ released versus time was plotted in the semi-logarithmic graph, and two linear regressions were fitted to natural logarithms of the rate of Mg^2+^ released. We interpreted that the first steep line represents the release of Mg^2+^ from the root surface and intercellular apoplastic spaces, and that the second line represents the Mg^2+^ release from the cytosol. The y-intercept of the second line represents the rate of Mg^2+^ release from cytosol at time zero, which indicates the efflux rate (pmol mg^-1^ min^-1^).

### Identifying the candidate gnomic region of LMGC1

An F2 progeny was developed by a cross between LMGC1 and wild type Koshihikari. The generated 90 individuals of F2 were grown with the control nutrient solution for 2 weeks, and the blade of 4th leaf was applied to the elemental analysis. Among the F2, DNA from 10 individuals having highest and lowest in Mg concentration were pooled to prepare High and Low Mg bulk samples, respectively. The bulked DNA samples were subjected to NGS and MutMap+ analysis (Fekih et al., 2013). After detecting the genomic region harboring contrast SNP-index values between High and Low Mg bulks, we compared the depth of aligned sequence reads between bulk samples. Finally, the genomic regions having few sequence reads specifically in Low Mg bulk sample were detected as candidate large deletion for phenotype of LMGC1. The association between this large deletion and Low Mg content was confirmed by other F2 progeny by PCR with the primer designed at flanking region of the detected deletion (Supplemental Table 1).

### Generation of knock-out mutants

#### Vector construction

The 20-nt annealed oligonucleotide pairs for the target sequences at the 1st exon of RZF (Os01g0555100) or 2nd exon of Rac (Os01g0555200) shown in the Supplemental Table 1 were cloned into the BbsI site of the single guide RNA (sgRNA) expression vector reported previously as pU6gRNA-oligo (Mikami et al., 2015). Two sgRNA expression cassettes targeting RZF (U6::sgRZF-1 and U6::sgRZF-2) or Rac (U6::sgRac-1 and U6::sgRac-2) were amplified by PCR with the primer sets (supplemental Table 1) and were tandemly combined and were inserted into AscI site in pZH_OsU6sgRNA _SpCas9 (Endo et al., 2019) by In-Fusion reaction (Takara), respectively. Finished CRISPR/Cas9 binary vectors were transformed into Agrobacterium tumefaciens EHA105 (Hood et al., 1993) by electroporation (E. coli Pulser, BioRad).

#### Transformation of rice callus

Agrobacterium-mediated transformation was performed as previously described (Toki, 1997). Rice (*Oryza sativa* L. cultivar Nipponbare) calli were grown on N6D medium at 33 °C for 4 weeks and were infected with Agrobacterium harboring the CRISPR/Cas9 binary vector. After 3 days of co-cultivation with Agrobacterium at 23 °C on solidified 2N6-AS medium, the calli were washed and transferred to N6D medium containing 25 mg/l meropenem (Fujifilm Wako Pure Chemical Industries) and 50 mg/l Hygromycin B (Fujifilm Wako Pure Chemical Industries) at 33 °C for 4 weeks. Hygromycin-resistant calli were subjected to HMA assay to identify the callus lines containing mutations at the target site. The callus lines containing mutations at the target site were transferred to a regeneration medium containing 25 mg/l meropenem and cultured at 28°C under 16 h light:8 h dark conditions for 4 weeks to obtain the regenerated mutant plants.

#### Heteroduplex mobility assay (HMA)

Genomic DNA was extracted from hygromycin-resistant calli using a one-step DNA extraction protocol (Kasajima et al., 2004). The RZF or Rac loci containing sgRNA target sites were amplified using KOD one PCR master mix (TOYOBO) and the primers listed in the Supplemental Table 1. PCR products were analyzed using MCE-202 MultiNA with a DNA-500 kit (SHIMADZU).

### Gene expression analysis

Rice cultivar Nipponbare was grown with control nutrient solution for 1 week, and the blade and the sheath of the third leaf as well as the root were sampled. RNA was extracted with the use of TRIsure reagent (Bioline), and 500 ng of which was used for reverse transcription using SuperScript IV (Invitrogen, USA). The relative amount of accumulation of *OsRZF1* (Os01g0555100) was determined using *OsGAPDH* gene as a control (Santos et al., 2018) by PCR method using KOD one PCR master mix (TOYOBO) (Supplemental Table 1).

### Protein localization assay

The ORF of OsRZF1 gene without stop codon was amplified (Supplemental Table 1) and cloned to the entry vector with pENTR™/D-TOPO^®^ Cloning Kit (Invitrogen, USA). Then the 2×35s promoter (Marquès-Bueno et al., 2016) as well as amplified OsRZF1 ORF were inserted into R4pGWB604 plasmid (Nakamura et al., 2010) using Gateway™ LR Clonase™ II Enzyme mix (Thermo Fisher Scientific, USA). The constructed expression vector was then transformed into *Agrobacterium tumefaciens* strain GV3101 competent cells by electroporation. Single colonies were picked and cultured overnight, washed, and resuspended in Agromix buffer containing 10 mM MES (pH 5.8), 10 mM MgCl2, and 150 μM acetosyringone. To perform transient expression, 8-week-old *Nicotiana benthamiana* leaves were inoculated by filtration through a 1-mL syringe with the resuspended solution. After 48 hours, the EGFP fluorescence was detected using FV-1200 confocal laser-scanning microscope (Olympus, Japan). Images were analyzed and merged by ImageJ (NIH).

### Statistical analysis

The data of growth parameters, Mg content, and ^28^Mg uptake were analyzed using unpaired Student’s t-test to compare the wild-type and the mutants. In the boxplots, the interquartile range with the median was presented. The mean value with standard deviation was presented in the bar chart and the scattergram.

## Acknowledgements and Funding

Authors thank Mrs. Midori Takemura for her assistance of plant cultivation. This work was supported by Japan Society for the Promotion of Science (Grant-in-Aid for Scientific Research (C) 19K05751 to N.I.K., Grant-in-Aid for Scientific Research (B) 20H02885 to K.T., and Grantin-Aid for Scientific Research (A) 20H00437 to T.M.N.).

---

**Supplemental Table 1.**
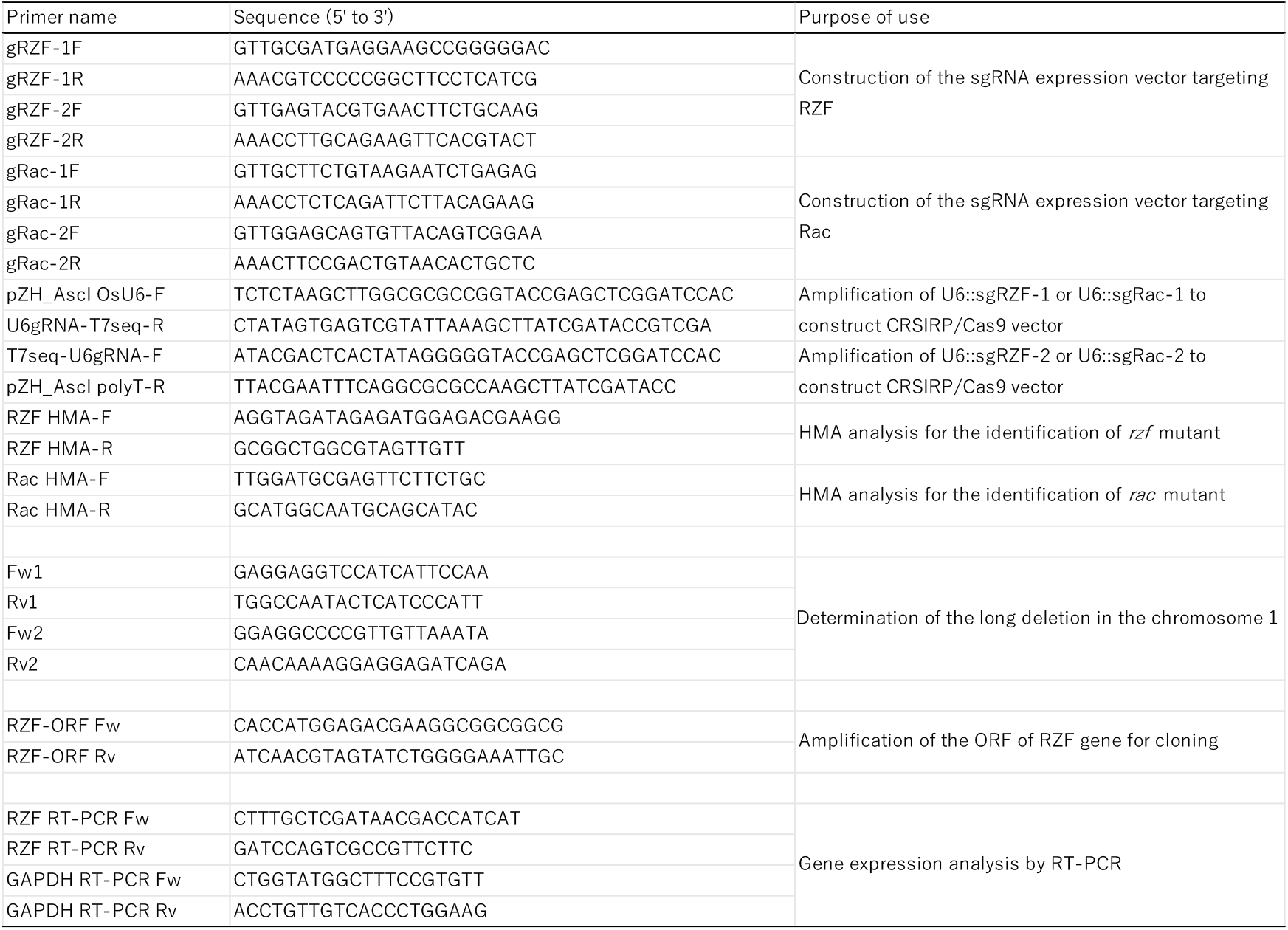
List of primers.

